# Label-free imaging of immune cell dynamics in the living retina using adaptive optics

**DOI:** 10.1101/2020.07.07.191890

**Authors:** Aby Joseph, Colin J Chu, Guanping Feng, Kosha Dholakia, Jesse Schallek

## Abstract

Our recent work characterized the movement of single blood cells within the retinal vasculature of healthy mice (Joseph et al. 2019) using adaptive optics ophthalmoscopy. Here we apply this technique to the context of acute inflammation and discover both infiltrating and tissue-resident immune cells to be visible without any labelling in the living retina using near - infrared light alone. Intravital imaging of immune cells can be negatively impacted by surgical manipulation, exogenous dyes, transgenic manipulation and phototoxicity. These confounds are now overcome, using phase contrast and time-lapse videography to reveal the dynamic behavior of myeloid cells as they interact, extravasate and survey the retina. Cellular motility and differential vascular responses can be measured noninvasively and in vivo across hours to months at the same retinal location, from initiation to the resolution of inflammation. As comparable systems are already available for clinical research, this approach could be readily translated to human application.

**Impact statement:** Immune cell motility and vascular response are imaged *in vivo* and label-free in the CNS for the first time, using high-resolution phase-contrast adaptive optics retinal imaging.

## Introduction

A robust and noninvasive method for directly imaging immune cells in humans has not been established, limiting our understanding of autoimmune and inflammatory disease. Immune cells typically provide weak optical contrast requiring exogenous or transgenic labelling for detection even in animal models. The eye is optimally suited to image immune responses *in vivo* [1]. As the only transparent organ in mammals, no inflammatory surgical or environmental perturbation is required. However, aberrations of the otherwise clear optics of the eye limit achievable image resolution. We employed a custom adaptive-optics-scanning-light-ophthalmoscope (AOSLO) to image the mouse eye. By correcting for the eye’s aberrations, single-cell resolution provides detailed imaging of photoreceptors to erythrocytes [2–4]. Incorporating our recent report [5] on phase contrast approaches allowed deep tissue detection of translucent cells [6,7]. We combined and applied these strategies and serendipitously discovered that immune cells could be imaged without labelling using 796 nm near-infrared light, to which the eye is insensitive, and at far lower levels (<500 µW) than multiphoton systems that can be phototoxic [8]. This new approach builds on our ability to quantify single-cell blood flow in vessels [2] revealing the dynamic interplay of blood flow and single immune cells in response to inflammation in the living eye.

## Results and Discussion

To model an immune response, ocular injection of lipopolysaccharide (LPS) was used to provide an acute but self-resolving inflammatory stimulus [9]. Potential immune cells were observed adjacent to retinal veins only after meticulous image registration, frame averaging and time-lapse imaging (**Fig. 1A, Video 1**). Membrane remodelling, pseudopodia formation and motility consistent with immune cells was visible, distinct from static neurons or macroglia (**Fig. 1B & Video 2)**. Within post-capillary venules, leukocyte rolling, crawling, and trans-endothelial migration behaviours were detectable (**Fig. 1C, Videos 3 and 4**). Heterogeneity in cell distribution, size and morphology was imaged with multiple cell types in different stages of interaction (**Fig. 1D**).

**Figure 1.**
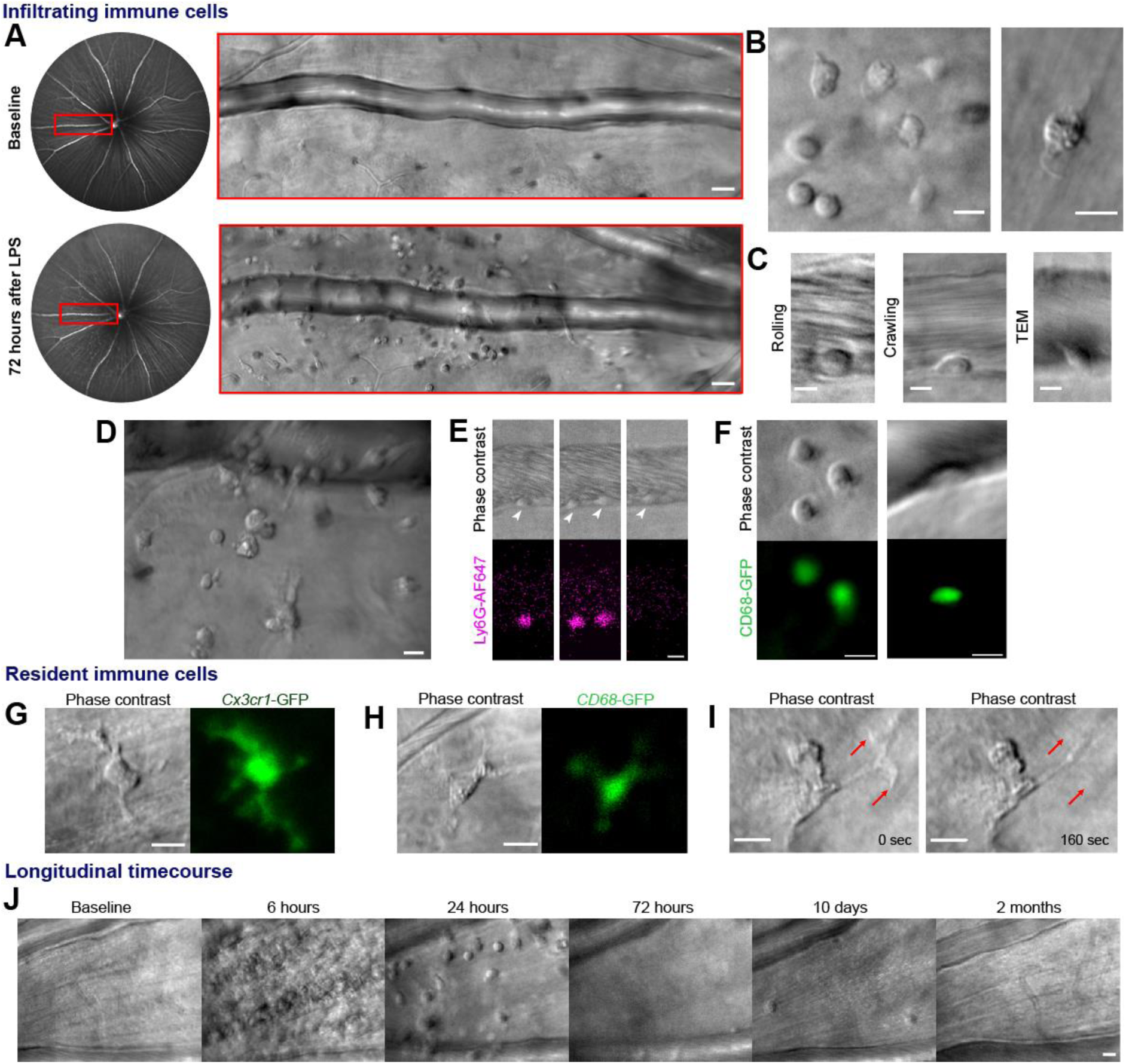
Label-free adaptive optics imaging of infiltrating and tissue resident immune cells in the retina. **A**, widefield image showing venule baseline and 72-hours after LPS injection. AOSLO montage (*red rectangle*) detects dispersed immune cells. **B**, detail of heterogenous immune cells **C**, Intravascular trans-endothelial migration (TEM) stages are visible. **D**, field adjacent to vein 72-hours after LPS. **E**, simultaneous phase-contrast and anti-Ly6G fluorescence reveals leukocyte rolling (*arrowheads*) and **F**, *CD68*-GFP reporter shows extravascular and intravascular cells. **G**, Representative examples of tissue-resident myeloid cells from *Cx3cr1*-GFP and **H**, *CD68*-GFP reporter mice showing colocalization of fluorescence with label-free cells. **I**, phase-contrast image of process remodelling. **J**, longitudinal imaging (hours-to-months) at same location following LPS injection. Scale bars =10 µm, except in **A** =50 µm.

We verified these cells comprised neutrophil and monocyte populations by fluorescent marker co-localization. Simultaneous phase contrast and confocal fluorescence AOSLO revealed most leukocytes rolling along venular endothelium were neutrophils using intravenous anti-Ly6G antibody labelling (**Fig. 1E, Video 5**). Conversely, *CD68*^*GFP/+*^ mice distinguished a population of cells were infiltrating monocytes and macrophages present both in vessels and extravasated into retinal tissue (**Fig. 1F**). More cells were visible using phase contrast than by fluorescence labelling, demonstrating its utility for comprehensively detecting diverse and mixed cellular populations. Tissue resident myeloid cells were also visible by AOSLO phase contrast even in healthy eyes without LPS injection. These were confirmed as microglia or hyalocytes by colocalization of *Cx3cr1*^*GFP/+*^ and *CD68*^*GFP/+*^ fluorescence (**Fig. 1G-I**) [10]. Phase contrast even revealed subcellular features, including structures that could represent internal processes such as endosomes (**Fig. 1H, Video 6**) [11].

As our approach is uniquely non-invasive, repeated imaging at the same tissue location permits longitudinal study throughout the initiation, peak and resolution of an immune response across hours to months within individual eyes (**Fig. 1J, Video 7**). To quantify immune cell behaviour in these studies, we had to distinguish immune cells from surrounding tissue by developing semi-automated deep learning software [12]. This correlated well with counts made by masked human observers (R^2^=0.99, P=0.004, **Video 8 (last part)**).

Immune cell metrics were quantified in six mice over five timepoints following LPS injection (**Fig. 2A-C**). Compared to baseline (28.9 ± 34.1 cells/mm^2^, Mean±SD) a 7-fold influx of cells was detected by six hours post injection (208.3 ± 108.6 cells/mm^2^) rising to over an 18-fold increase by 24 hours (510.4 ± 441 cells/mm^2^) before returning towards baseline at 72-hours (59.0 ± 27.8 cells/mm^2^) and 10-days (69.4 ± 41.7 cells/mm^2^). AOSLO also allowed cell motility quantification with maximum cell displacement observed at 6 hours (16.1 ± 9.9 µm, n=12 cells). Despite peak infiltration at 24-hours, motility was greatly reduced (4.6 ± 4.3 µm, n = 58 cells), best appreciable by longitudinal imaging, consistent with Resolvin-mediated suppression of chemotaxis [13].

**Figure 2.**
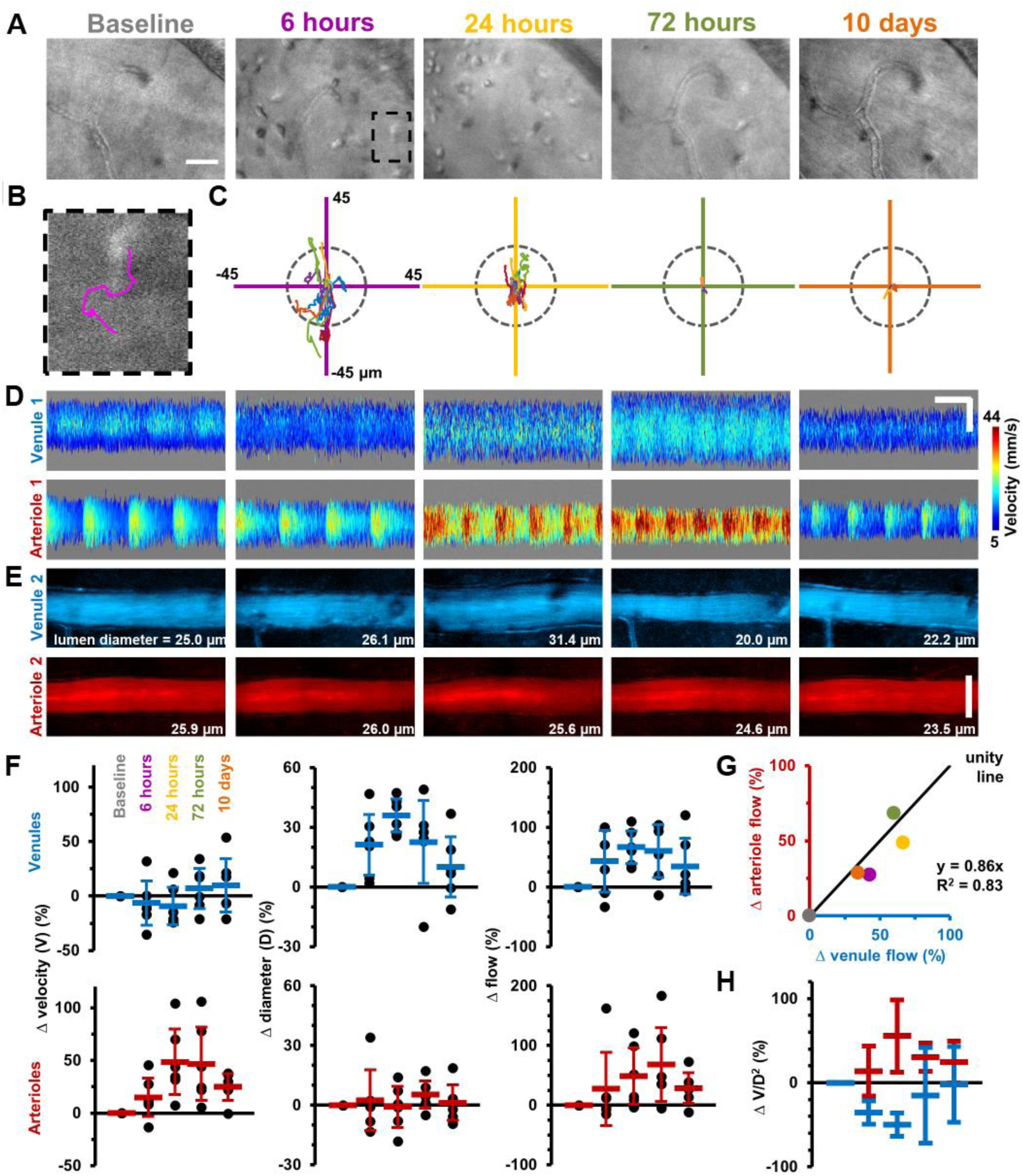
Longitudinal non-invasive measurement of combined immune cell dynamics and vascular flow. Measurements of neighbouring venules, arterioles and connecting parenchyma. **A**, AOSLO phase-contrast images of same region across five timepoints relative to LPS injection. Scale bar =30 µm. **B**, Magnified image of one cell (marked overlay) with tracked trace (100 seconds). **C**, Cell displacements for total cohort at indicated timepoints. Displacement traces normalized to each cell starting position. Grey dashes indicate radius of typical cell size (13 µm). **D**, Space-time images with overlaid single-cell blood velocity. Arteriolar velocity increases then resolves. Scale bars=200 ms horizontal, 30 µm vertical. **E**, Vessel diameter visualized in motion-contrast images of venules (blue) and arterioles (red). Venule diameter dilates then resolves. Scale bar=30 µm **F**, Population values of RBC velocity change relative to baseline, lumen diameter and flow rate for venules and arterioles. Vein diameter (p=0.008) and artery velocity (p=0.036) exhibit significant changes across time (Friedman test, n=6 mice). Mean+SD shown. **G**, Correlation of change in flow between arterioles and venules, (colors correspond to timepoints in **A**). **H**, Change in ratio V/D^2^, plotted across timepoints from **F** (V=velocity, D=diameter). Flow-rate is proportional to the product V*D^2^.

As retinal tissue is not depressurized by this intravital system, true vascular alterations arising from inflammation can be isolated and correlated to simultaneous immune cell measurements. Adapting our recent work [2], red blood cell (RBC) velocimetry, vessel dilation and flow-rate changes were quantified in this same cohort of mice (**Fig. 2D-H**). AOSLO revealed micron-level vascular dilations and heterogeneous changes in blood flow in arterioles and venules in response to LPS. Total blood flow increased in the retinal circulation, yet elevated flow in arterioles and venules was achieved in fundamentally different ways. Venules dilated on average 36% (±8%) at 24-hours post LPS injection facilitating a total flow increase of 67% (±27%) relative to baseline. Conversely, arterioles showed minimal dilation yet dramatically elevated RBC velocity (48% increase (±31%) at 24-hours (**Fig. 2D-F**). Despite these different mechanisms, conservation of flow was confirmed as arterioles and venules showed correlated changes in flow over time (R^2^=0.83, for linear-fit, **Fig. 2G-H**). Velocity, dilation and flow began resolution towards baseline levels by 10-days post-injection. The balanced asymmetry between arterioles and venules is a new finding in our understanding of hemodynamic control in response to inflammation, achieved by applying our recently developed tool [2].

This study advances our understanding of the immune response by imaging both the nuanced differences of the vascular perfusion with the first detailed imaging of single immune cell activity imaged without labels. Unlike other approaches, inflammatory surgery is not required, the label-free feature avoids confounding by exogenous fluorescent dyes, gene haplo-insufficiency from transgenic labels and using near-infrared light avoids phototoxicity inherent to multiphoton platforms [8,14]. By providing this proof of concept, as AOSLO systems are already available for clinical research the approach stands to be rapidly translated to human application.

## METHODS

### Mouse strains

All mice were sourced from The Jackson Laboratory (Bar Harbor, Maine, USA) and maintained at the University of Rochester in compliance with all guidelines from the University Committee on Animal Resources and according to the Association for Research in Vision and Ophthalmology statement for the Use of Animals in Ophthalmic and Vision Research. Mice were fed with standard laboratory chow *ad libitum* and housed under a 12-hour light-dark cycle. 6 to 12-week old male mice were employed from three strains: C57BL/6J (Stock number 000664), *hCD68-GFP* (Stock number 026827) and *Cx3cr1*-GFP hemizygotes (Stock number 005582).

### Mouse preparation for imaging

Mice underwent anesthetic induction with intraperitoneal Ketamine (100 mg/kg) and Xylazine (10 mg/kg) before maintenance on 1% v/v isofluorane and supplemental oxygen through a nose cone. Pupils were dilated with a single drop of 1% Tropicamide and 2.5% phenylephrine (Akorn, Lake Forest, IL, USA). Internal temperature was controlled using an external heating pad adjusted to maintain continuous 37.0 degrees Celsius with monitoring via a rectal probe electrical thermometer (Physiosuite, Kent). A rigid contact lens of 1.6mm base curve and +10 Dioptre correction was placed centrally on the cornea and lubrication of the eye maintained by aqueous lubricant (GenTeal, Alcon Laboratories, Fort Worth, TX, USA) during imaging. The eye was imaged in free space meaning there was no physical contact with the AOSLO, ensuring no compression causing alteration of intraocular pressure.

### Endotoxin induced uveitis model

Following anesthesia, intravitreal injection with a 34-gauge Hamilton microsyringe through the pars plana was used to deliver 0.5 ng of lipopolysaccharide (LPS) from *E.coli* 055:B5 (Sigma) in a one microlitre volume of phosphate buffered saline (PBS) [9]. Only one eye of each mouse was injected and used for the study.

### Intravenous antibody labelling

2 µg of primary conjugated anti-mouse Ly6G-Alexa Fluor 647 (clone 1A8, Biolegend) diluted into 200 µl PBS were injected intravenously via tail vein ten minutes prior to imaging as previously published [15,16].

### AOSLO imaging

Mice were imaged with a custom adaptive optics scanning light ophthalmoscope (AOSLO), using near-infrared light (796Δ17 nm, 200-500 µW, super luminescent diode: S790-G-I-15, Superlum, Ireland) [2,17]. Phase-contrast imaging referred to in the context of this paper, was achieved by purposefully displacing the detector axially to a plane conjugate to the highly reflective RPE/choroid complex, to enable detection of forward and multiply scattered light from translucent cells, as detailed in our recent publication [5]. In a subset of experiments for confirmation of immune cell types, fluorescence was simultaneously imaged using 488 nm excitation and 520Δ35 emission for GFP, and 640 nm excitation and 676Δ29 emission for Alexa Fluor 647 (excitation laser diode: iChrome MLE, Toptica Photonics, Farmington, New York, USA; emission filters: FF01-520/35-25 and FF01-676/29-25, Semrock, Rochester, New York, USA). Mice also underwent imaging with HRA+OCT Spectralis (Heidelberg Engineering, Germany).

### Aberration measurement and correction with adaptive optics

Aberrations in the mouse eye were measured and corrected with a closed-loop adaptive optics system operating at 13 corrections per second, built at the University of Rochester and described previously [2,17]. Aberrations were measured with a Hartmann-Shack wavefront sensor using a 904 nm wavefront beacon (QFLD-905–10S, QPhotonics, Ann Arbor, Michigan, USA) imaged onto the retina. Aberration correction was achieved with a continuous membrane deformable mirror with 97 actuators (DM-97-15, ALPAO, France).

### Imaging videography

The AOSLO is a raster-scanning instrument with a resonant scanner frequency of 15 kHz and 25 Hz orthogonal scanning rendering the retina at 25 frames per second. Point scanning readout was achieved by two photomultiplier tubes (PMTs) for visible and near infrared wavelengths (H7422–40 and H7422–50, Hamamatsu, Japan). Frame size was 608 by 480 pixels and the image distortion introduced by sinusoidal scanning was corrected in real-time [18]. Field sizes were between 2-5 degrees of visual angle corresponding to 68-170 microns in retinal space. Typical imaging sessions lasted ∼2 hours and semi-continuous video acquisition of a target retinal location was conducted for up to 30 minutes. AOSLO retinal data was visualized in real-time to facilitate user tracking, correction and optimization and saved for subsequent post-processing.

### Image registration and time-lapse analysis

To correct for residual motion of the eye, image registration was performed with custom software [18,19]. Time-lapse videos were generated using running frame-averaging of 25-50 frames (from 25 Hz native frame rate of AOSLO) with ImageJ and Fiji (National Institutes of Health, USA) [20]. Montaging multiple fields in Fig. 1a was performed by stitching and blending overlapping 4.98 degree fields manually using Adobe Photoshop (Version: CS6 Extended v13.0.1 x64).

### Statistical analysis

Pearson correlation and Freidman tests were performed using Prism 7.0 (GraphPad Software) and linear curve-fitting in MATLAB 2020a, version 9.8.0.

### Cell migration measurement

Retinal locations were imaged for 100 seconds for six mice at five timepoints (baseline, 6, 24, 72-hours and 10-days post LPS injection). Sixty AOSLO phase-contrast videos were used to analyse the migration of extravasated cells. Registered videos were pre-processed by cropping to 512×400 pixels and temporally averaged with five frames. A customized semi-automated deep-learning based cell tracking software was employed to track and quantify the migration behaviour of cells. The software consisted of a deep learning-based cell detector with an encoder-decoder U-Net backbone architecture [12,21]. The U-Net was trained on phase-contrast AOSLO images with the centroids of 387 cells manually identified by expert graders obtained from four mice at either 6 or 24-hours post LPS injection. Immune cell data from mice imaged for training were excluded from the final analysis of the results in this paper. The trained U-Net outputs a probability map for each frame of a video that was thresholded (>= 90%) to identify the centroids of cells. The cell counts of a video were calculated by averaging the number of detected objects in the first 25 frames (5 seconds). To track the cells, centroid positions detected by the U-Net in adjacent frames were linked with a nearest neighbor search algorithm [21]. The deep learning strategy facilitated tracking of a large number of cells across multiple frames captured at different time points across inflammation. Once the heavy burden of tracking this population over a large data set was complete, a human user provided quality control of the automated outputted traces to confirm tracking fidelity. Traces due to incorrect linkage (for example, a single trace that falsely jumped between two adjacent cells) were manually rejected based on visual inspection (Fig. 2b, Video 8). To quantify cell migration, two quantities were extracted from the traces: 1. cell displacement which is defined as the displacement of a cell over 100 seconds. 2. confinement ratio which is defined as the ratio of cell displacement and the total path length over 100 seconds. Traces in Fig. 2c indicate the total positional movement and direction of movement relative to cell position in the first second of data collection. The U-Net based cell detection was performed with Python 3.7 and PyTorch 1.0.1, while the tracking and quantification procedures were based on MATLAB.

### Blood flow measurement

Single cell blood flow was imaged and measured with near infrared light using our recently published approach [2], for the same six mice and five timepoints as above, in an arteriole and venule surrounding the tissue location at which the cell migration measurements above were done. Briefly, a fast 15 kHz beam (796Δ17 nm) was scanned across a vessel of interest to image passing blood cells without requiring contrast agent. Cellular-scale blood velocity and vessel diameter were quantified automatically. Given the small size of even the largest mouse retinal vessels (<45 µm inner diameter in healthy mice), the spatial resolution of our approach accurately measured the inner lumen diameter with micrometer precision, accounting for vessel tortuosity, vascular wall thickness and cell-free plasma layer. Additionally, the temporal resolution of the velocity detection approach was more than sufficient to measure and account for cardiac pulsatility in flow, as demonstrated previously. This ensured accurate measurement of the average blood velocity through the vessel. Combined, the volumetric flow rate through the vessel was quantified label-free. The non-invasive approach enabled us to track blood flow longitudinally from hours to weeks over the course of inflammation without requiring invasive injections or euthanasia after a single timepoint was imaged.

### Code availability statement

Single-cell blood flow was measured using our recently published approach [2], with custom code written in MATLAB R2017a (Version 9.2, with Image Processing Toolbox, MathWorks, Massachusetts, USA). Source code is available in public repository here: https://github.com/abyjoseph1991/single_cell_blood_flow. Other code used in the study is available from corresponding authors upon reasonable request.

## Acknowledgments

The authors wish to thank Qiang Yang, Andres Guevara, Rachel Hollar, Jennifer Strazzeri and Karteek Kunala for their technical contributions to this work. We are grateful to Andrew Dick and Richard Lee for their guidance and helping establish this collaboration. We thank David Williams and Robin Sharma for their critical feedback on the manuscript.

## Supporting information: video legends

**Video 1 (separate file).** Demonstration of image post-processing. a. Raw adaptive optics scanning light ophthalmoscope (AOSLO) corrected 796nm phase contrast imaging of C57BL/6J mouse retina. Real-time video obtained at 25 fps acquisition (labelled “raw acquisition”) demonstrating movement from respiration and cardiac output. Image following custom frame-registration. Application of 25 frame temporal averaging and accelerated time-lapse (labelled “frame averaging”). Top row, cluster of infiltrated immune cells 6 hours post-LPS. Bottom row, tissue resident cell in healthy retina adjacent to a retinal capillary. Scale bars = 10 μm.

**Video 2 (separate file).** Label-free AOSLO time-lapse video demonstrating heterogenous immune cell populations and motility. Two magnified locations from Fig. 1b showing 796nm phase contrast video acquired at 25 frames per second from two C57BL/6J mouse retinas 24 hours post-LPS injection. The second segment of the video is a full 4.98-degree AOSLO field at a retinal vein 48-hours after LPS injection, revealing a diversity of cell morphologies and motility patterns. Videos have undergone post-processing as described, 25-50 frame temporal averaging. Scale bars = 10 μm.

**Video 3 (separate file).** Leukocyte rolling and crawling in an inflamed retinal post-capillary venule. 796nm phase contrast AOSLO images taken at 6 hours post-LPS injection with 25 frame temporal averaging. Descriptive overlays provided. Scale bar = 10 μm.

**Video 4 (separate file).** Examples of diverse immune cell behaviour observable by AOSLO phase contrast imaging. 796nm reflectance AOSLO images at 6 or 24 hours post-LPS using between 5 to 50 frame averaging. Examples include post-capillary venule leukocyte rolling, transendothelial migration and perivascular leukocyte accumulation, venous leukocyte rolling and crawling with and against blood flow direction, mid tissue infiltrating leukocyte swarming, perivascular cell process contact with intravascular cell and cell migration towards lumen of retinal vein. Scale bars = 10 μm.

**Video 5 (separate file).** Neutrophil endothelial rolling within a post-capillary venule confirmed using fluorescent labelling with anti-Ly6G antibody. Representative example of retinal vein imaged six hours post LPS injection. Simultaneous aligned acquisition of 796nm phase contrast (top panel) and anti-Ly6G conjugated AlexaFluor 647 (positively labelling neutrophils in bottom panel) using confocal AOSLO fluorescence. Scale bar = 10 μm.

**Video 6 (separate file).** Cx3cr1-GFP+ tissue resident cell with motile processes. Simultaneous aligned acquisition of 796nm phase contrast (left panel) and GFP fluorescence (right panel) in healthy retina from Cx3cr1GFP/+ reporter mouse. Second section demonstrates process retraction in tissue resident myeloid cell. Third section illustrates diversity of morphology observed. Forth section highlights visible sub-cellular features in this population. Scale bar = 10 μm.

**Video 7 (separate file).** Repeated longitudinal imaging of the same retinal location from initial inflammation to resolution. Representative recordings from one C57BL/6J mouse at the same anatomical location in the retina, identifiable by peripapillary location and capillary and vascular landmarks. Recorded prior to and 6, 24, 72 hours, 10 days and 2 months following a single LPS injection. Scale bar = 10 μm.

**Video 8 (separate file).** U-Net cell tracking trace example. Recording from C57BL/6J mouse retina at 6 hours post LPS injection. Whole field imaged and quantified is shown on the left side. Magnification of one representative cell on the right. Scale bar = 16 μm. Trace is marked with an overlaid magenta line. Second section shows validation of U-Net cell

